# The effect of stimulus intensity on neural envelope tracking

**DOI:** 10.1101/2020.08.11.245761

**Authors:** Eline Verschueren, Jonas Vanthornhout, Tom Francart

**Author notes:** Correspondence: Eline Verschueren Herestraat 49, bus 721, 3000 Leuven, Belgium.

## Abstract

**Objectives:** The last years there has been significant interest in attempting to recover the temporal envelope of a speech signal from the neural response to investigate neural speech processing. The research focus is now broadening from neural speech processing in normal-hearing listeners towards hearing-impaired listeners. When testing hearing-impaired listeners speech has to be amplified to resemble the effect of a hearing aid and compensate peripheral hearing loss. Until today, it is not known with certainty how or if neural speech tracking is influenced by sound amplification. As these higher intensities could influence the outcome, we investigated the influence of stimulus intensity on neural speech tracking.

**Design:** We recorded the electroencephalogram (EEG) of 20 normal-hearing participants while they listened to a narrated story. The story was presented at intensities from 10 to 80 dB A. To investigate the brain responses, we analyzed neural tracking of the speech envelope by reconstructing the envelope from EEG using a linear decoder and by correlating the reconstructed with the actual envelope. We investigated the delta (0.5-4 Hz) and the theta (4-8 Hz) band for each intensity. We also investigated the latencies and amplitudes of the responses in more detail using temporal response functions which are the estimated linear response functions between the stimulus envelope and the EEG.

**Results:** Neural envelope tracking is dependent on stimulus intensity in both the TRF and envelope reconstruction analysis. However, provided that the decoder is applied on data of the same stimulus intensity as it was trained on, envelope reconstruction is robust to stimulus intensity. In addition, neural envelope tracking in the delta (but not theta) band seems to relate to speech intelligibility. Similar to the linear decoder analysis, TRF amplitudes and latencies are dependent on stimulus intensity: The amplitude of peak 1 (30-50 ms) increases and the latency of peak 2 (140-160 ms) decreases with increasing stimulus intensity.

**Conclusion:** Although brain responses are influenced by stimulus intensity, neural envelope tracking is robust to stimulus intensity when using the same intensity to test and train the decoder. Therefore we can assume that intensity is not a confound when testing hearing-impaired participants with amplified speech using the linear decoder approach. In addition, neural envelope tracking in the delta band appears to be correlated with speech intelligibility, showing the potential of neural envelope tracking as an objective measure of speech intelligibility.

## 1 INTRODUCTION

The last years there has been significant interest in attempting to recover the temporal envelope of a speech signal from the neural response to investigate neural speech processing (e.g., Ding and Simon, 2012b; O’Sullivan et al., 2015; Das et al., 2016). The temporal envelope contains syllable, word and sentence boundaries and is considered to be essential for speech understanding (Shannon et al., 1995). A growing body of research shows that the quality of reconstruction of the temporal envelope from the neural responses, so-called neural envelope tracking, is related to speech intelligibility (e.g. Ding and Simon, 2013; Di Liberto et al., 2018; Vanthornhout et al., 2018; Lesenfants et al., 2019; Decruy et al., 2019; Iotzov and Parra, 2019; Verschueren et al., 2020): The better the acoustic speech envelope is encoded in the brain, the better the speech is understood.

The research focus is now broadening from neural speech processing in normal-hearing listeners towards hearing-impaired listeners. Several studies already compared normal-hearing to hearing-impaired listeners: Petersen et al. (2017) and Presacco et al. (2019) reported no significant difference in neural envelope tracking of the attended speech between normal-hearing and hearing-impaired listeners. Decruy et al. (2020b), on the other hand, showed increased neural envelope tracking in hearing-impaired listeners compared to normal-hearing peers. The results of Mirkovic et al. (2019) are in line with Decruy et al. (2020b) and suggest (no statistics were reported) higher envelope tracking for the attended speech in participants with hearing loss. An important remark is that in three (Petersen et al., 2017; Mirkovic et al., 2019; Decruy et al., 2020b) out of four studies speech was (linearly) amplified. Amplification was applied to the stimuli to resemble the effect of a hearing aid and compensate peripheral hearing loss. It is not known with certainty how or if neural envelope tracking is influenced by amplification. Therefore it is difficult to disentangle whether the reported results above are an effect of hearing loss, or an effect of sound amplification. In addition, Verschueren et al. (2019) reported increased neural envelope tracking in the delta band with increasing speech understanding by varying the intensity of the presented speech in cochlear implant (CI) users. Again, it is difficult to distinguish whether neural envelope tracking decreases because of decreasing speech intelligibility or stimulus intensity.

In this study we investigate the influence of stimulus intensity on neural envelope tracking in normal-hearing listeners. We hypothesize that neural envelope tracking is sensitive to the absolute level at which the speech is presented. The link between stimulus intensity and cortical auditory evoked potentials (CAEP) has namely been thoroughly investigated and established using non-speech stimuli. In short, the amplitudes of the CAEP peaks increase and the latencies decrease with increasing stimulus intensity (e.g., Moore and Rose, 1969; Picton et al., 1970; Adler and Adler, 1989; Billings et al., 2007). In addition, a similar effect of stimulus intensity on brainstem responses has been shown using transient stimuli. With decreasing stimulus intensity, the latencies of the auditory brainstem response (ABR) peaks increase and the amplitudes of the response peaks decrease until the hearing threshold is reached and no response is present anymore (Hall, 2007). In line with this, Picton et al. (2003) and Van Eeckhoutte et al. (2016) reported a similar decreasing amplitude pattern with decreasing stimulus intensity using auditory steady-state responses (ASSR).

In summary, all these studies show an effect of stimulus intensity on auditory evoked potentials in normal-hearing listeners. However, none of them used running speech as a stimulus. Only one study, to our knowledge, explored the effect of variations in stimulus intensity on neural speech tracking using running speech: Ding and Simon (2012a) presented speech in the presence of a competing talker. They kept the competing talker intensity constant, whereas the target speech was attenuated or amplified by maximally 8 dB. They reported no significant difference in neural envelope tracking for these small variations in stimulus intensity. In this study we increase the range of stimulus intensities used by Ding and Simon (2012a) to investigate whether their results can be extended to a wider range of stimulus intensities. This could shed some light on the effects reported in hearing-impaired listeners: Did Decruy et al. (2020b) and Mirkovic et al. (2019) find enhanced neural envelope tracking in hearing-impaired participants listening to amplified sound because of increased stimulus intensity or because of a different factor? Was the increase of neural envelope tracking with increasing stimulus intensity shown by Verschueren et al. (2019) in CI users driven by stimulus intensity or speech understanding?

## 2 MATERIAL AND METHODS

### 2.1 Participants

Twenty participants aged between 19 and 24 years (4 men and 16 women) took part in the experiment after providing informed consent. Participants had Dutch as their mother tongue and were all normal-hearing, confirmed with pure tone audiometry (thresholds ≤ 25 dB HL at all octave frequencies from 125 Hz to 8 kHz). The study was approved by the Medical Ethics Committee UZ Leuven / Research (KU Leuven) with reference S57102.

### 2.2 EEG experiment

Recordings were made in a soundproof and electromagnetically shielded room. Speech was presented bilaterally using APEX 4.0 (**?**), an RME Multiface II sound card (Haimhausen, Germany) and Etymotic ER-3A insert phones (Illinois, USA). The setup was calibrated using a 2 cm^3^ coupler of the artificial ear (Brüel & Kjær 4152, Denmark). EEG was recorded with a 64-channel BioSemi ActiveTwo EEG recording system was at a sample rate of 8192 Hz. Participants sat in a comfortable chair and were asked to move as little as possible during the recordings.

The story used during the EEG measurement was ‘The Casual Vacancy’ by J.K. Rowling, narrated in Dutch by Nelleke Noordervliet. The story was split in 6 longer blocks of ≈ 5 minutes [mean = 316 seconds, sd = 2.8 seconds] and 8 shorter blocks of ≈ 2 minutes [mean = 124 seconds, sd = 2.6 seconds] which were presented in chronological order. The 5-minute blocks were presented randomly at 30, 60 and 70 dB A, applying every intensity twice to obtain 10 minutes of speech at a same intensity, which was necessary for further analysis. The short 2-minute blocks were presented at 8 different intensities randomly selected from a fixed list, containing 10, 20, 30, 40, 50, 60, 70, 80 dB A. An overview of the different conditions is shown in Figure 1. Two out of 20 participants, participating in the pilot study, did not listen to speech intensities higher than 60 dB A. To ensure understanding of the story line, the story summary was shown before the experiment started. To maximize the participants attention and motivation, content questions were asked after each 2- or 5-minute block. In addition, speech intelligibility was measured after each block using a rating method where the participants were asked to rate their speech understanding on a scale from 0 to 100% following the question ‘Which percentage of the story did you understand?’.

**Figure 1.**
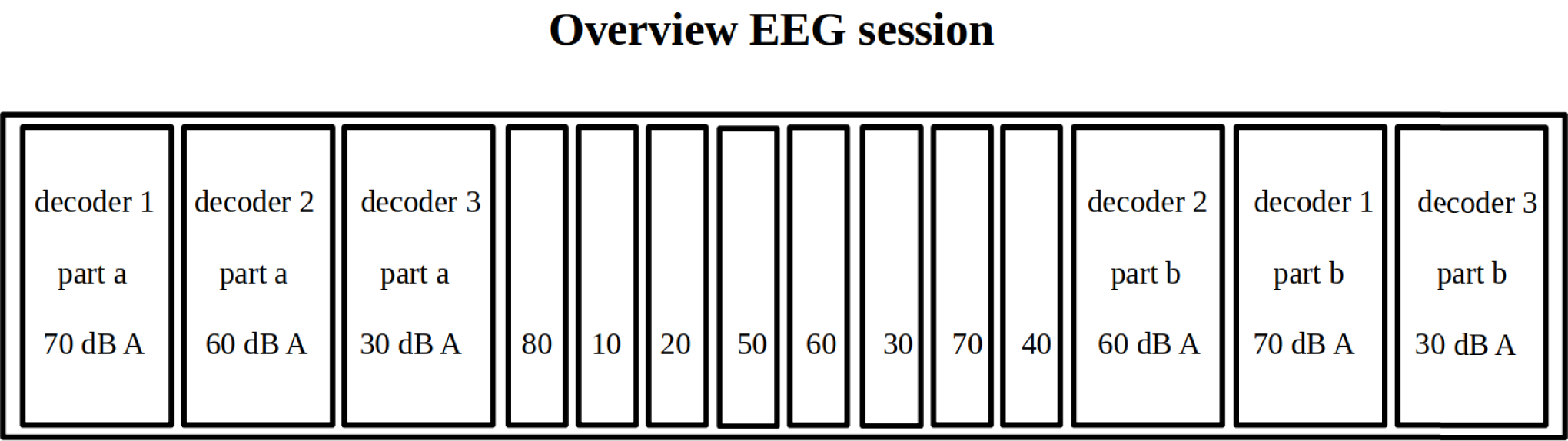
Overview of an example test session. The numbers in each block indicate the presented stimulus intensity in dB A. Six longer 5-minute blocks were presented at 30, 60 and 70 dB A. Every intensity was used twice (part a and b) to obtain 10 minutes of speech at a same intensity. Eight shorter blocks were presented at intensities varying from 10 to 80 dB A.

### 2.3 Signal processing

In this study we measured neural envelope tracking and related this to stimulus intensity. Neural envelope tracking was calculated using an envelope reconstruction approach (2.3.1) and temporal response functions (2.3.2).

#### 2.3.1 Envelope reconstruction

Details of the signal processing steps applied on the EEG data to measure neural envelope tracking using envelope reconstruction are described carefully in Vanthornhout et al. (2018) and Verschueren et al. (2020)).

In short, we extracted the speech envelope from the original signal according to Biesmans et al. (2017) using a gamma-tone filter bank followed by a power law, and band-pass filtered the envelope in the delta (0.5-4 Hz) and theta (4-8 Hz) frequency band. Second, we reconstructed the speech envelope from the EEG with a linear decoder estimated using ridge regression similar to Lalor et al. (2006), taking all 64 EEG channels and their time-lagged variants from 0 to 250 ms into account. In summary the following steps were applied: the EEG was downsampled from 8192 Hz to 256 Hz, artifact rejection was applied using the multi-channel Wiener filter (Somers et al., 2018), the signal was band-pass filtered similar to the acoustic speech envelope and the sample rate was further decreased from 256 Hz to 128 Hz. Only the longer 10-minute blocks at 30, 60 and 70 dB A were used to train the decoder. The shorter 2-minute blocks with varying intensity from 10 to 80 dB A were used to apply the decoder on. As a consequence the decoder could be applied on data of the same or a different intensity as it was trained on.

To investigate training and testing on a same stimulus intensity we applied leave-one-out cross-validation, after normalization, on the longer 10-minute speech blocks (Figure 2, upper panel). We always used 1 minute for testing and the remaining 9 minutes for training, resulting in 30 (3 stimulus intensities x 10 cross-validations) reconstructed envelopes per participant. To investigate training and testing on a different intensity we trained 2 decoders on the entire 10-minute blocks at 30 and 60 dB A. These decoders were used to reconstruct the envelope in the shorter 2-minute story blocks, after normalization, resulting in 16 (2 decoders x 8 stimulus intensities) envelopes per participant (Figure 2, lower panel).

**Figure 2.**
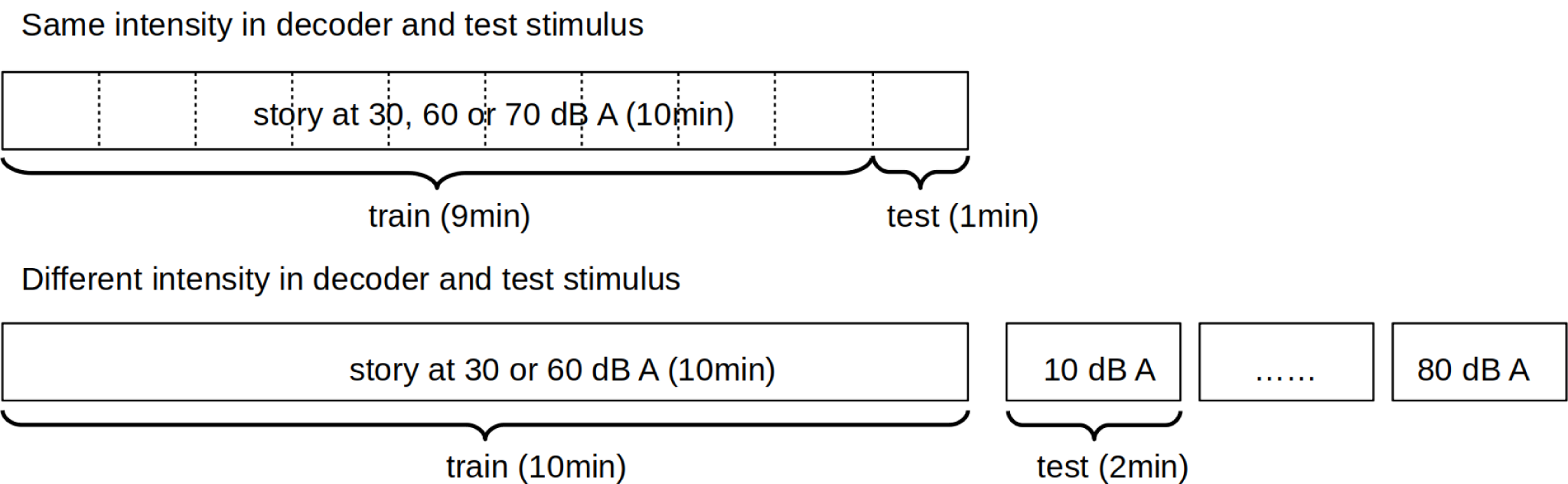
Overview of the 2 analysis methods used to reconstruct the envelope from the EEG response. The upper panel shows the leave-one-out cross-validation on the longer 10-minute speech blocks where decoder and test stimulus have the same intensity. The lower panel shows the decoder approach where a decoder is trained on a separate stimulus (10-minute block) and applied on the 2-minute blocks.

Finally, to quantify neural envelope tracking, we correlated the speech envelope obtained from this reconstruction analysis with the actual speech envelope using bootstrapped Spearman correlations. The significance level of the correlation was calculated by correlating random permutations of the real and reconstructed envelope 1000 times and taking percentile 2.5 and 97.5 to obtain a 95% confidence interval.

#### 2.3.2 Temporal response function estimation

The main advantage of the envelope reconstruction analysis is the integration of all EEG channels and time-lagged variants leading to a powerful prediction. This integration advantage is at the same time also a disadvantage: no interpretation is possible per electrode channel and per latency (Haufe et al., 2014). Therefore temporal response function (TRF) estimations are a valuable additional analysis giving some insights at a channel- and latency-level. In addition, no model assumptions are made like with decoders. A TRF is a simplified brain model that describes how the brain, if it was a linear filter, would process the acoustic speech envelope of the stimulus to obtain the recorded EEG signal.

We calculated a TRF for every electrode channel in every participant at 30, 60 and 70 dB A. We only included the longer 10-minute blocks because of the amount of data needed to obtain a good prediction. The first signal processing steps are identical to the envelope reconstruction model, only the filtering was done within a broader band (0.5-25 Hz). Next, TRFs were calculated using the boosting algorithm using the Eelbrain source code (Broderick et al., 2018) as described in detail by David et al. (2007). In summary, boosting is an iterative algorithm using the mean-squared error (MSE) to find the most optimal prediction for the given data. After calculation, the TRFs were convolved with a rotationally symmetric Gaussian kernel of 5 samples long (SD=2) to smooth over time lags.

### 2.4 Statistical Analysis

Statistical analysis was performed using MATLAB (version R2018a) and R (version 3.4.4) software. The significance level was set at α =0.05 unless otherwise stated.

To investigate the effect of stimulus intensity on neural envelope tracking we created a linear mixed effect model (LME) per filter band with the following general formula:

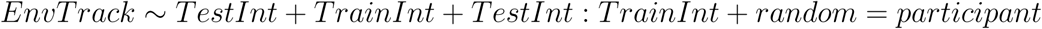

where “EnvTrack” refers to the Spearman correlation between the reconstructed and the acoustic envelope and “TestInt” refers to the intensity the test stimulus was presented at. We added a fixed and interaction effect of “TrainInt”, which is the intensity on which the decoder was trained. In addition, a random effect of intercept of the participants was included in the model to allow neural tracking to be higher or lower per participant. To control if every chosen fixed and random effect benefited the model the Akaike Information Criterion (AIC) was calculated. Unstandardized regression coefficients (beta) with 95% confidence intervals and p-value are reported in the results section.

Furthermore, we compared dependent samples using a non-parametric Wilcoxon signed-rank test. To compare multiple dependent samples with incomplete observations we used the Skillings–Mack statistic (Skillings et al., 1981), an extension of Friedman’s test (e.g. neural envelope tracking at 30, 60 and 70 dB A: not all participants listened to 70 dB A speech as mentioned in section 2.2).

## 3 RESULTS

We analyzed behavioral speech intelligibility in function of stimulus intensity (section 3.1. Next, we investigated the neural responses to the speech signal at different stimulus intensities in two ways. First we reconstructed the envelope from the EEG using a decoder and correlated this reconstructed envelope with the actual acoustic envelope (section 3.2). Second, we calculated TRFs to investigate the neural response per channel over time (section 3.3).

### 3.1 Behavioral speech intelligibility

To investigate the effect of stimulus intensity on speech intelligibility we asked the participants to rate their speech intelligibility after each presented 2-min block of the story. Figure 3 shows that speech intelligibility increases with increasing stimulus intensity until reaching a plateau at ≈ 30 dB A. Fitting a sigmoid function on the data shows that participants understand 50% of the presented speech signal at 17.6 dB A (midpoint of the sigmoid function). Inter-subject variability depends on the intensity conditions (Levene’s test, F=7.76, p<0.001) with the largest variability at an intensity of 20 dB A (sd=22.42).

**Figure 3.**
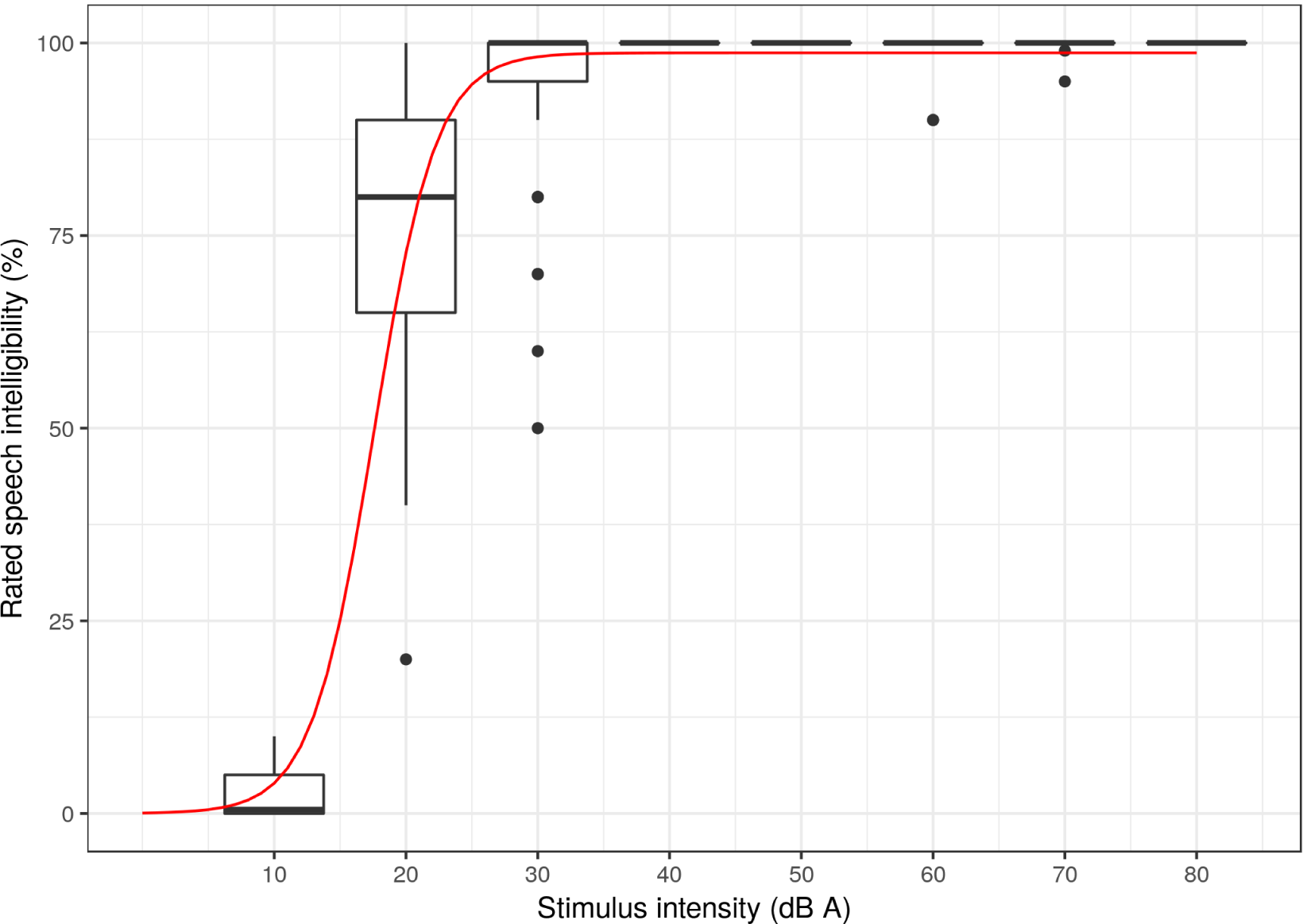
Rated speech intelligibility in function of stimulus intensity. Rated speech intelligibility increases with increasing stimulus intensity, saturating at 30 dB A. Variability between participants is largest in the 20 dB A condition. The red line shows the psychometric function fitted on the data.

### 3.2 Effect of stimulus intensity on envelope reconstruction

#### 3.2.1 Training and testing on the same stimulus intensity

Figure 4 shows the results of the leave-one-out cross-validation on the longer 10-minute speech blocks. The train and test stimulus were presented at the same stimulus intensity. No significant difference was found in neural envelope tracking between a stimulus presented at 30, 60 or 70 dB A in the delta (p=0.55, Skillings-Mack statistic = 1.20) or theta band (p=0.52, Skillings-Mack statistic = 1.30).

**Figure 4.**
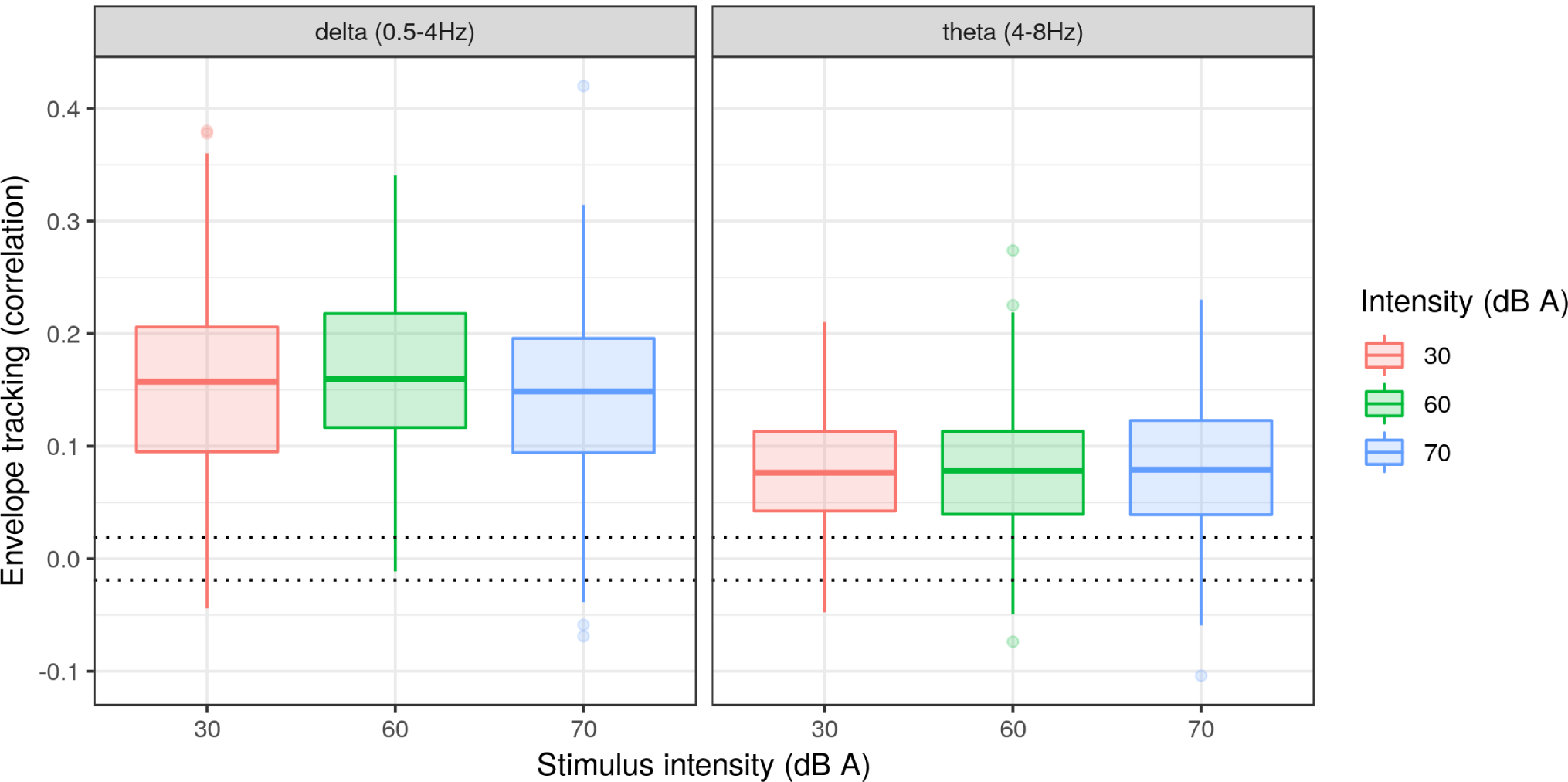
Envelope tracking in function of stimulus intensity when using the same intensity to train and test the decoder. No significant difference in envelope tracking was found between a stimulus presented at 30, 60 and 70 dB A. The dotted lines show the significance level of the correlation.

#### 3.2.2 Training and testing on a different stimulus intensity

For the next analysis, we trained a decoder for the delta and theta band at 30 and 60 dB A, using the longer 10-minute blocks of the story. These decoders were applied on the shorter 2-minute blocks with varying intensities from 10 to 80 dB A, resulting in 32 envelopes (2 decoders x 2 filter bands x 8 stimulus intensities) per participant.

Figure 5 shows an increase of neural envelope tracking with increasing stimulus intensity in the delta band (p<0.001, b=0.0005, CI(95%): ±0.0002, LME). However, when excluding the 10 dB A (median speech intelligibility = 2.6%) condition from statistical analysis and keeping all other remaining conditions with a speech intelligibility level of 50% or more, no significant effect of stimulus intensity on neural envelope tracking can be found (p=0.2, b=-0.0002, CI(95%): ±0.0003, LME). The stimulus intensity on which the decoder was trained does not add extra information to the model in the delta band (LME, AIC = −874.06 versus AIC = −874.63).

**Figure 5.**
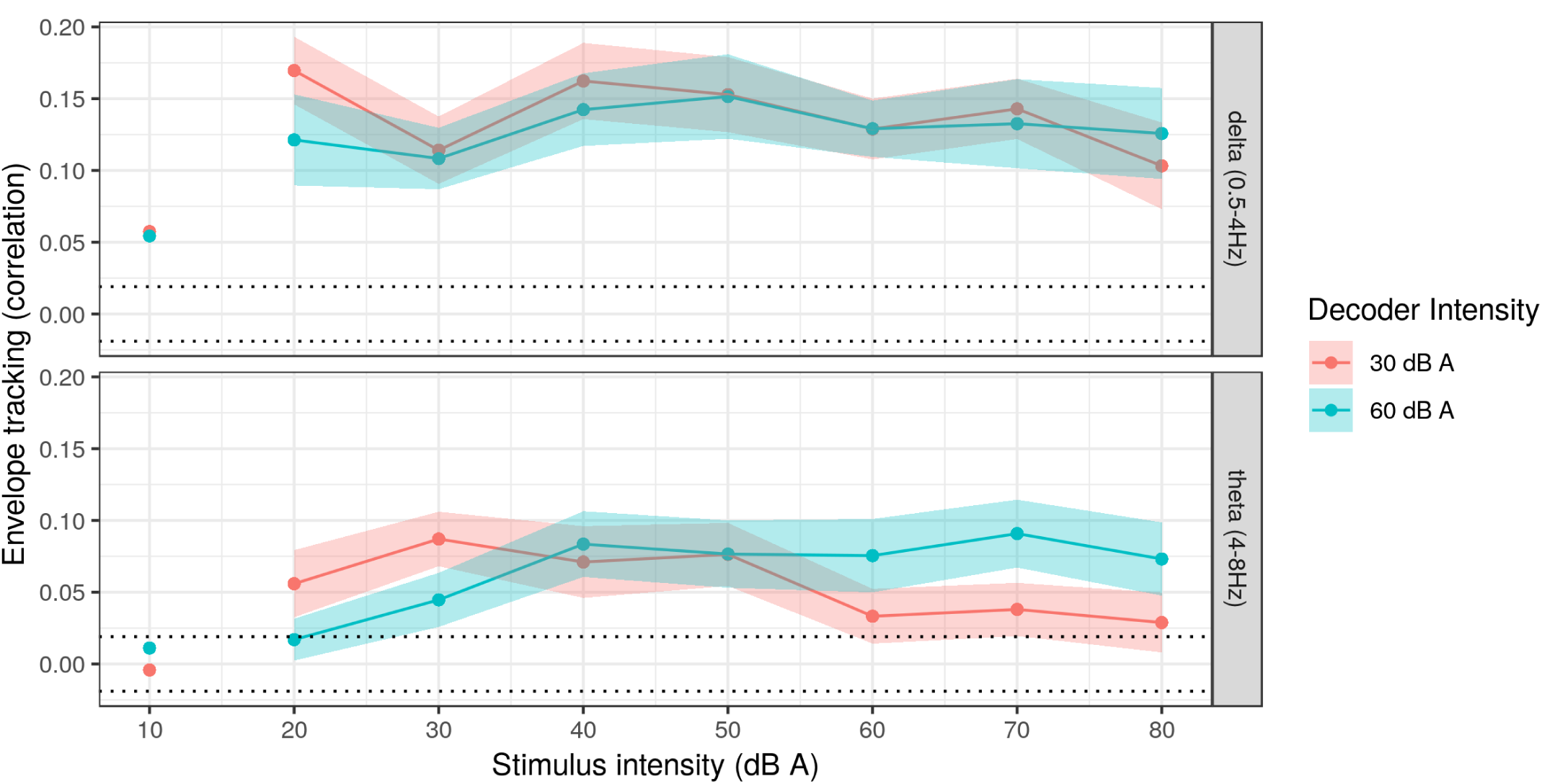
Envelope tracking in function of stimulus intensity when using a different intensity to train and test the decoder. Envelope tracking is not influenced by stimulus intensity in the delta band. In the theta band tracking increases with increasing stimulus intensity for a 60 dB A decoder, but decreases for a 30 dB A decoder. The solid line shows the median over subjects, the shading is 2 times the standard error. The dotted lines show the significance level of the correlation.

In the theta band, in contrast to the delta band, the effect of stimulus intensity on neural envelope tracking (p<0.001, b=0.0006, CI(95%): ±0.0002, LME) does not disappear when removing the 10 dB A condition (p<0.001, b=-0.0008, CI(95%): ±0.0004, LME). Moreover, the stimulus intensity on which the decoder was trained influences the absolute level of neural envelope tracking (p<0.001, b=-0.0732, CI(95%): ±0.0273, LME) and the slope of neural envelope tracking over stimulus intensity (p<0.001, b=0.0017, CI(95%): ±0.0005, LME). Figure 5 shows that, in the theta band, neural envelope tracking increases with increasing stimulus intensity for the 60 dB A decoder (slope LME = 0.96 x 10^−3^). For the 30 dB A decoder, on the other hand, neural envelope tracking decreases with increasing stimulus intensity (slope LME = −0.75 x 10^−3^). More in detail, a Wilcoxon signed rank test reveals that the envelope of 60 dB A speech can be reconstructed with higher accuracy using a decoder trained at 60 dB A compared to a decoder trained at 30 dB A (p=0.008, CI(95%) = [-0.072; −0.027], n=20, Wilcoxon signed-rank test). Similarly, the envelope of 30 dB A speech can be reconstructed with higher accuracy using a decoder trained at 30 dB A compared to a decoder trained at 60 dB A (p<0.001, CI(95%) = [0.012; 0.055], n=20, Wilcoxon signed-rank test). In summary, a decoder trained on a low intensity improves envelope reconstruction at low intensities, while a decoder trained on a high intensity improves envelope reconstruction at high intensities.

#### 3.2.3 Effect of speech intelligibility on envelope reconstruction

Besides comparing stimulus intensity to neural envelope tracking, it is also of interest to analyze the potential relation between actual speech intelligibility and neural envelope tracking. To investigate this, we selected the condition with the most inter-subject variability: intensity = 20 dB A (analysis in section 3.1). Within this condition, speech intelligibility varied from 20 to 100% between participants while stimulus intensity remained constant. Based on the previous results showing that best envelope reconstructions are obtained when using a decoder trained on the same or a similar intensity, we selected the 30 dB A decoder to reconstruct the envelope at 20 dB A (There was not sufficient data to train a decoder at 20 dB A). Figure 6 shows that neural envelope tracking increases with increasing speech intelligibility in the delta band (r=0.57, p=0.01, Spearman rank correlation): The better a participant can understand very soft speech, the higher neural envelope tracking for that participant. In the theta band no significant effect was found (r=0.00, p=0.99, Spearman rank correlation).

**Figure 6.**
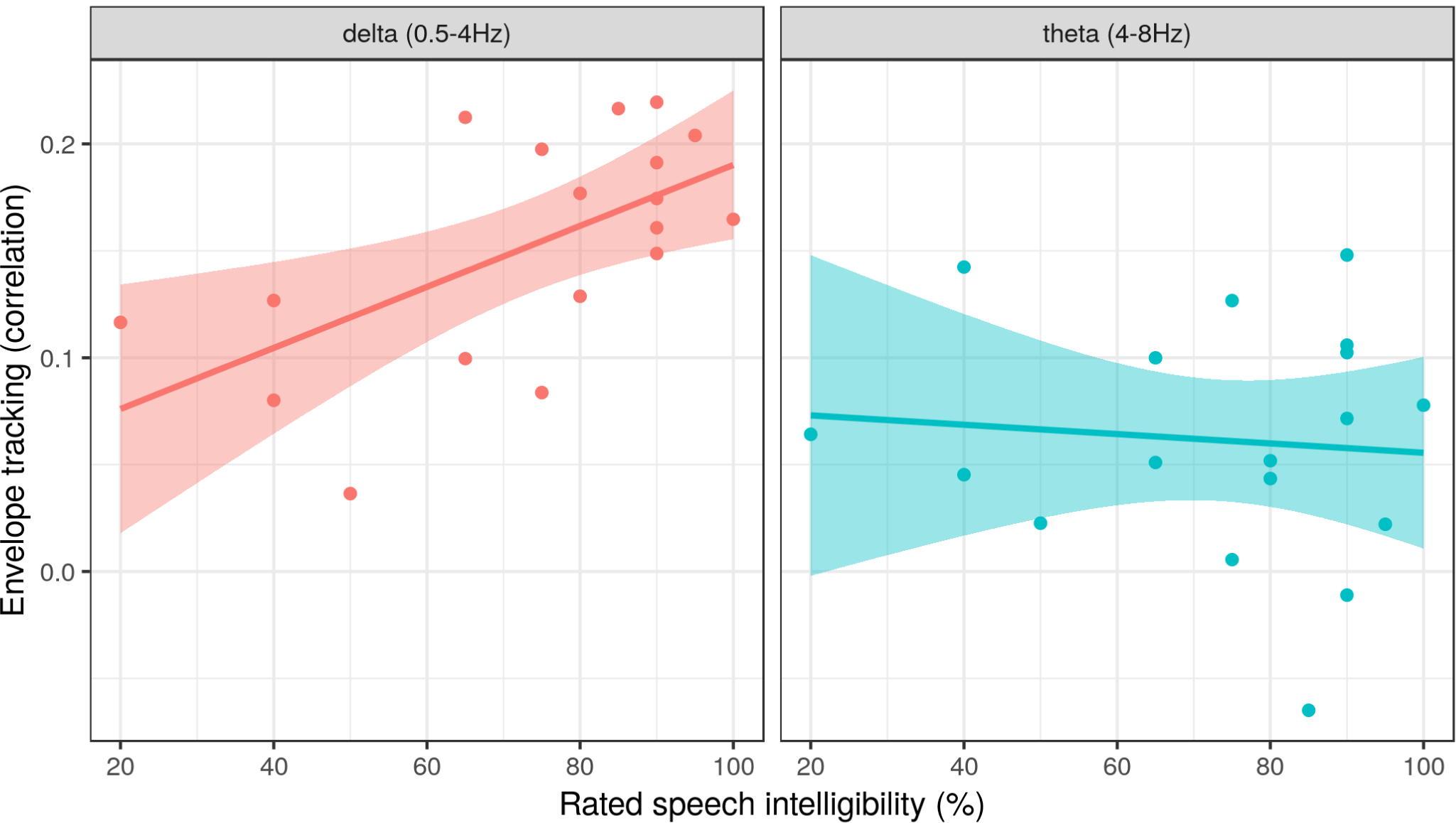
Envelope tracking increases with increasing speech intelligibility at a stimulus intensity of 20 dB A in the delta band, but not in the theta band. Every dot represents a participant. The solid line shows the linear model fit per filter band with the 95% confidence interval in shading.

### 3.3 Effect of stimulus intensity on TRFs

The envelope reconstruction analysis showed that neural envelope tracking is not significantly influenced by stimulus intensity when using the same intensity to train and test the decoder. However, when training and testing on a different stimulus intensity, the quality of envelope reconstruction in the theta band depends on how similar the two intensities are. To better understand where this stimulus-induced difference originates from, we created TRFs of every 10-minute segment (30, 60, 70 dB A). Based on visual inspection of the topographies we selected the centro-frontal channels as the channels of interest (Figure 7, black dots on panel A.).

**Figure 7.**
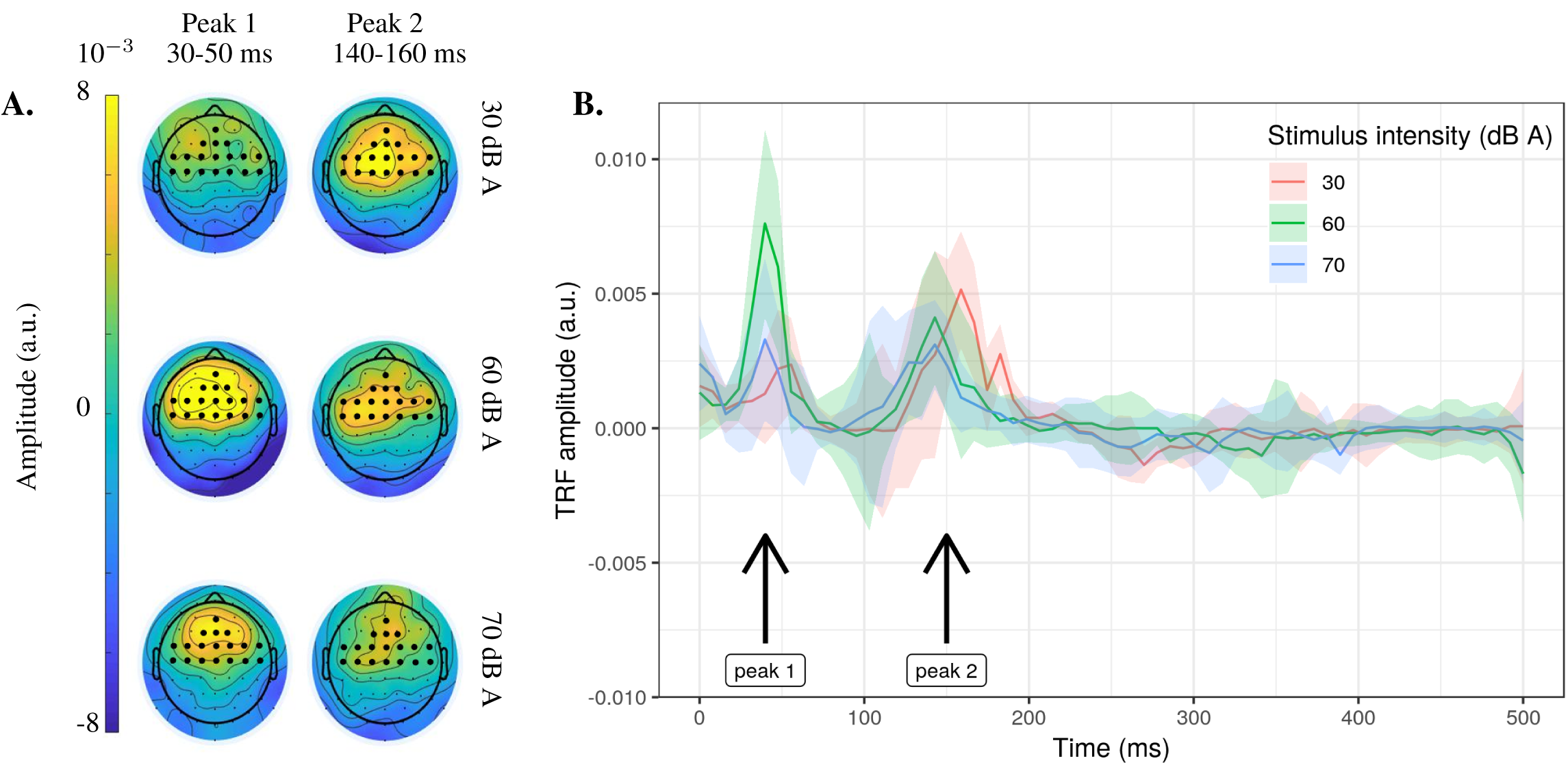
Temporal response functions (TRF) per stimulus intensity. A. Topographies showing median TRF activity over subjects at the latencies of peak 1 and 2. Black dots represent the centro-frontal channel selection to create the TRFs in time-domain in panel B. B. Median centro-frontal TRF over participants per intensity, shading is the standard error over participants.

Figure 7B shows 2 prominent peaks around 30-50 ms and 140-160 ms. Figure 7A shows the topographies of both peaks. To investigate the influence of stimulus intensity on the latency and amplitude of these peaks, we calculated the maximum for every participant per stimulus intensity between respectively 20 and 70 ms, and 100 and 200 ms (= the peak value). The latency of the first peak is not significantly affected by stimulus intensity (p=0.55, Skillings-Mack statistic = 1.20). The amplitude, on the other hand, is affected by stimulus intensity (p=0.01, Skillings-Mack statistic = 9.30): When stimulus intensity increases from 30 to 60 dB A, the amplitude of peak 1 increases (p=0.002, α=0.025, CI(95%) = [-0.010; −0.003], n=20, Wilcoxon signed-rank test). However, when stimulus intensity further increases to 70 dB A, peak 1 decreases as shown in Figure 7B (p=0.004, α=0.025, CI(95%) = [0.001; 0.007], n=18, Wilcoxon signed-rank test). Peak 2, in contrast to peak 1, shows no significant effect of stimulus intensity on amplitude (p=0.212, Skillings-Mack statistic = 3.10), but on peak latency (p=0.036, Skillings-Mack statistic = 6.65): When stimulus intensity increases from 30 to 60 dB A the latency of peak 2 decreases (p=0.003, α=0.025, CI(95%) = [3.97; 23.81], n=20, Wilcoxon signed-rank test). When stimulus intensity further increases to 70 dB A latency saturates (p=0.37, α=0.025, CI(95%) = [-23.81; 7.94], n=18, Wilcoxon signed-rank test), as shown in Figure 7B.

## 4 DISCUSSION

We investigated whether stimulus intensity influences neural envelope tracking. We tested 20 normal-hearing participants who listened to a story at different stimulus intensities. Neural envelope tracking is dependent on stimulus intensity in both TRF and envelope reconstruction analysis. However, provided that the decoder is applied on data of the same stimulus intensity as it was trained, envelope reconstruction is robust to stimulus intensity. This is promising for applications in hearing-impaired listeners requiring amplified speech. In addition, neural envelope tracking in the delta (but not theta) band seems to relate to speech intelligibility.

### 4.1 Effect of stimulus intensity on envelope reconstruction

#### 4.1.1 Robust envelope reconstruction when training and testing on the same stimulus intensity

We found no significant difference in neural envelope tracking between 30, 60 or 70 dB A when the decoder was applied on data of the same stimulus intensity as it was trained. This is consistent with the results reported by Ding and Simon (2012a) where no significant difference in neural envelope tracking was found when speech was attenuated or amplified by maximally 8 dB. We can thus tentatively conclude that presenting speech at different intensity levels, to for example ensure speech intelligibility in hearing-impaired listeners, does not influence the results. This implies that the enhancement of neural envelope tracking in hearing-impaired compared to normal-hearing listeners reported by Mirkovic et al. (2019) and Decruy et al. (2020b) is not driven by sound amplification. A plausible explanation for their enhanced results could be that hearing-impaired listeners need more effort to perform in the same way as normal-hearing listeners. Increased listening effort, and thus increased attention, has been associated with enhanced neural responses (Decruy et al., 2020a; Vanthornhout et al., 2019; Ding and Simon, 2012a,b).

#### 4.1.2 Training and testing on different stimulus intensities affects neural envelope tracking

Besides training and testing on a same stimulus intensity as mentioned in Section 4.1.1, we also trained and tested on different stimulus intensities. When using different stimulus intensities to train and test the decoder on, reconstruction accuracy changes, especially in the theta band: a decoder trained on a low intensity enhances envelope reconstruction at low stimulus intensities, while a decoder trained on a high intensity enhances envelope reconstruction at high stimulus intensities. The brain responses differ in function of stimulus intensity, as also shown in the TRF analysis 3.3. The more the intensities for the training and testing part are similar (ideally equal), the smaller the difference in brain responses and the better the reconstruction accuracy. This underlines the importance of using a decoder stimulus that is very similar to the testing stimulus for optimal tracking accuracy.

### 4.2 Neural envelope tracking in the delta (but not theta) band relates to speech intelligibility

Next, we compared neural envelope tracking results to speech intelligibility. We hypothesized that the better the acoustic speech envelope is encoded in the brain, the better the speech is understood (Ding and Simon, 2013; Di Liberto et al., 2018; Vanthornhout et al., 2018; Verschueren et al., 2020; Lesenfants et al., 2019; Decruy et al., 2019). In the delta band, stimulus intensity did not influence neural envelope tracking significantly between 20 (median SI = 80%) and 80 dB A (median SI = 100%). Only when including the lowest condition of 10 dB A (median SI = 2.6%) an increase of neural envelope tracking with increasing stimulus intensity could be found. This suggests that delta band tracking correlates with speech intelligibility rather than stimulus intensity. Only when speech intelligibility drops, neural envelope tracking in the delta band drops. In the theta band, on the other hand, an effect of stimulus intensity can be found even though speech intelligibility remains constant. This frequency band discrepancy between the delta and theta band could be explained by the hypothesis proposed by Ding and Simon (2014). They state that the delta band is related to the prosody of speech, while the theta band encodes syllabic-level acoustic features. Perhaps prosodic features are encoded more robustly in the brain compared to syllabic features, making them less sensitive to stimulus intensity variations while speech intelligibility remains the same.

An additional argument to strengthen the hypothesis of speech intelligibility versus acoustics in respectively the delta and the theta band are the results in Figure 6: Neural envelope tracking increases with increasing speech intelligibility in the delta band, but not in the theta band. This is in line with results reported by Molinaro and Lizarazu (2017). They also found that the theta band reflects perceptual processing of the auditory signal, while the delta band involves additional higher order speech specific processing. In addition, studies by Meyer et al. (2017) and Vanthornhout et al. (2018) also show that especially the delta band tracking correlates with respectively syntactic processing and speech intelligibility. Moreover, these results showing that delta band tracking is relatively robust to stimulus intensity and could reflect speech intelligibility clarify the results reported by Verschueren et al. (2019): The increase in neural envelope tracking with increasing stimulus intensity in cochlear implant users could be related to speech intelligibility rather than stimulus intensity.

### 4.3 Stimulus intensity affects temporal response functions

When stimulus intensity increased from 30 to 60 dB A, the amplitude of peak 1 (30-50 ms) increased. This is in line with the observed increase of auditory evoked potential amplitudes in function of stimulus intensity for non-speech stimuli (Moore and Rose, 1969; Picton et al., 1970; Adler and Adler, 1989; Billings et al., 2007; Hall, 2007; Picton et al., 2003; Van Eeckhoutte et al., 2016). However, when further increasing the stimulus intensity to 70 dB A, the amplitude of peak 1 decreased. A possible explanation for this non-linear behavior of TRF amplitude over stimulus intensity could be the saturation of the neural response. Several studies have reported amplitude saturation and decrease at around 70 dB (Butler et al., 1969; Picton et al., 1970; Adler and Adler, 1989), which is the same stimulus intensity as reported in this study.

In contrast to peak 1, the amplitude of peak 2 (140-160 ms) did not depend on stimulus intensity. This difference between peak 1 and 2 could be explained by the hypothesis that the early latencies of the TRF reflect acoustic processing of the speech (peak 1), while the later latencies (peak 2) of the TRF reflect higher-order speech processing (e.g., phonetic processing). An intensity-dependent peak 1 amplitude is thus to be expected, as a change in intensity is an acoustical change. A change in peak 2 amplitude, on the other hand, is not to be expected because a change in stimulus intensity does not influence speech understanding and thus higher-order speech processing. Ding and Simon (2013) reported similar results using speech in background noise. When more noise was added (the quality of the speech signal decreased), the amplitude of the TRF peak at 50 ms decreased (~peak 1), while the amplitude of the 100 ms peak (~peak 2) remained unchanged as long as speech could be understood.

Besides the amplitude, also the latency of the TRF peaks can be analyzed. The latency of peak 1 was not significantly affected by stimulus intensity. The latency of peak 2, on the other hand, was affected by stimulus intensity: When stimulus intensity decreased, the latency of peak 2 increased from 30 to 60 dB A. We hypothesize that the TRF peak is delayed with lower stimulus intensities because, although behaviorally measured speech intelligibility remains the same, the lower stimulus intensity makes it harder to understand the speech, resulting in a longer time needed to process the speech. This is similar to auditory evoked response potential literature using non-speech stimuli where decreasing stimulus intensities result in increasing latencies (Hall, 2007; Picton et al., 2003; Van Eeckhoutte et al., 2016).

### 4.4 Implications for applied research

In this study we showed that neural envelope tracking depends on stimulus intensity for both the TRF and envelope reconstruction analysis. However, when the same stimulus intensity is used to train and test the decoder, envelope reconstruction seems not to be affected by stimulus intensity in contrast to TRFs. The apparent discrepancy between envelope reconstruction and TRFs is due to the underlying model: as the brain response function changes with intensity (as apparent from our TRF analysis), intensity-specific decoders are required. However, when this is the case, the brain response is similarly strong across intensities, as long as speech can be understood.

An interesting question that arises from this apparent discrepancy is which method (TRF vs decoder) is most optimal to measure envelope tracking. We believe both methods have their advantages and disadvantages, depending on the intended purpose. TRFs can provide us with a lot more spatial and temporal information about the participants speech processing. In addition this method appears sensitive enough to find differences in acoustic properties of the speech the participant is listening to (e.g., stimulus intensity). The downside is that interpreting TRFs is time-consuming and can only be done by highly trained personnel. Decoders, on the other hand, are much more robust. Because of this robustness to for example stimulus intensity decoders seem a reliable method when testing hearing-impaired participants with amplified speech and comparing them to normal-hearing peers. In addition, decoders do not require any channel selection or peak-picking and return one correlation, making it easy to interpret.

When considering the application of envelope tracking as an objective measure of speech intelligibility both methods could be complimentary. Decoders could for example be used to screen which patient has decreased speech intelligibility and should be referred to an audiologist. After referral, TRFs could for example be used to diagnose the specific speech intelligibility problem and decide on an adequate treatment.

### 4.5 Conclusion

In this study we showed that neural envelope tracking is dependent on stimulus intensity in both TRF and envelope reconstruction analysis. However, provided that the decoder is applied on data of the same stimulus intensity as it was trained on, envelope reconstruction is robust to stimulus intensity. Therefore we can assume that intensity is not a confound when testing hearing-impaired participants with amplified speech using the linear decoder approach. Second, we showed that higher envelope tracking can be obtained in the theta band when the intensity to train and test the decoder are similar. In addition, neural envelope tracking in the delta band appears to be correlated with speech intelligibility, showing the potential of neural envelope tracking as an objective measure of speech intelligibility.

## Acknowledgements

The authors would like to thank Lien Decruy, Sofie Keunen and Elise Verwaerde for their help in data acquisition.

